## Supplementary Figures for "Copy number variation introduced by a massive mobile element underpins global thermal adaptation in a fungal wheat pathogen"

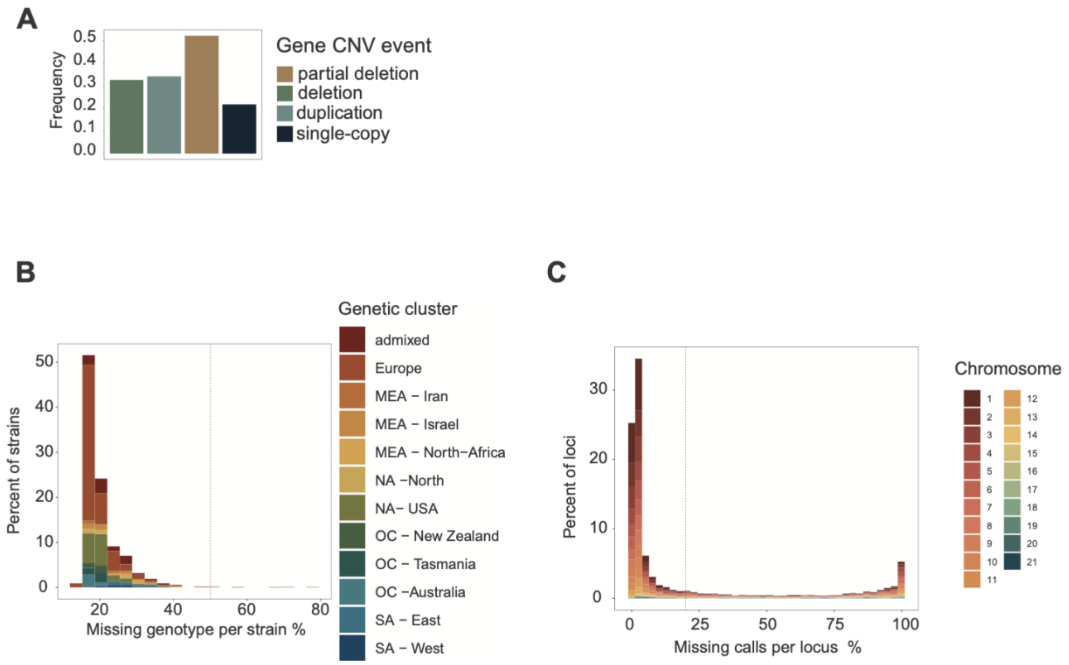


**Supplementary Figure 1.** A) Structure variant call comparison between gene CNV calls in the global collection (based on GATK) and chromosome-level assemblies (based on SyRI) for matching four isolates. B) Frequency of gene CNV event calls removed after filtering on CNQ scores in the global collection (GATK pipeline) and validation by the chromosome-level assemblies (SyRI analyses). C) Distribution of missing genotypes per strain in the CNV call dataset. The red dotted line refers to the 50% threshold. D) Distribution of missing calls per locus (*i.e.* gene) in the CNV call dataset. The red dotted line refers to the 20% threshold.

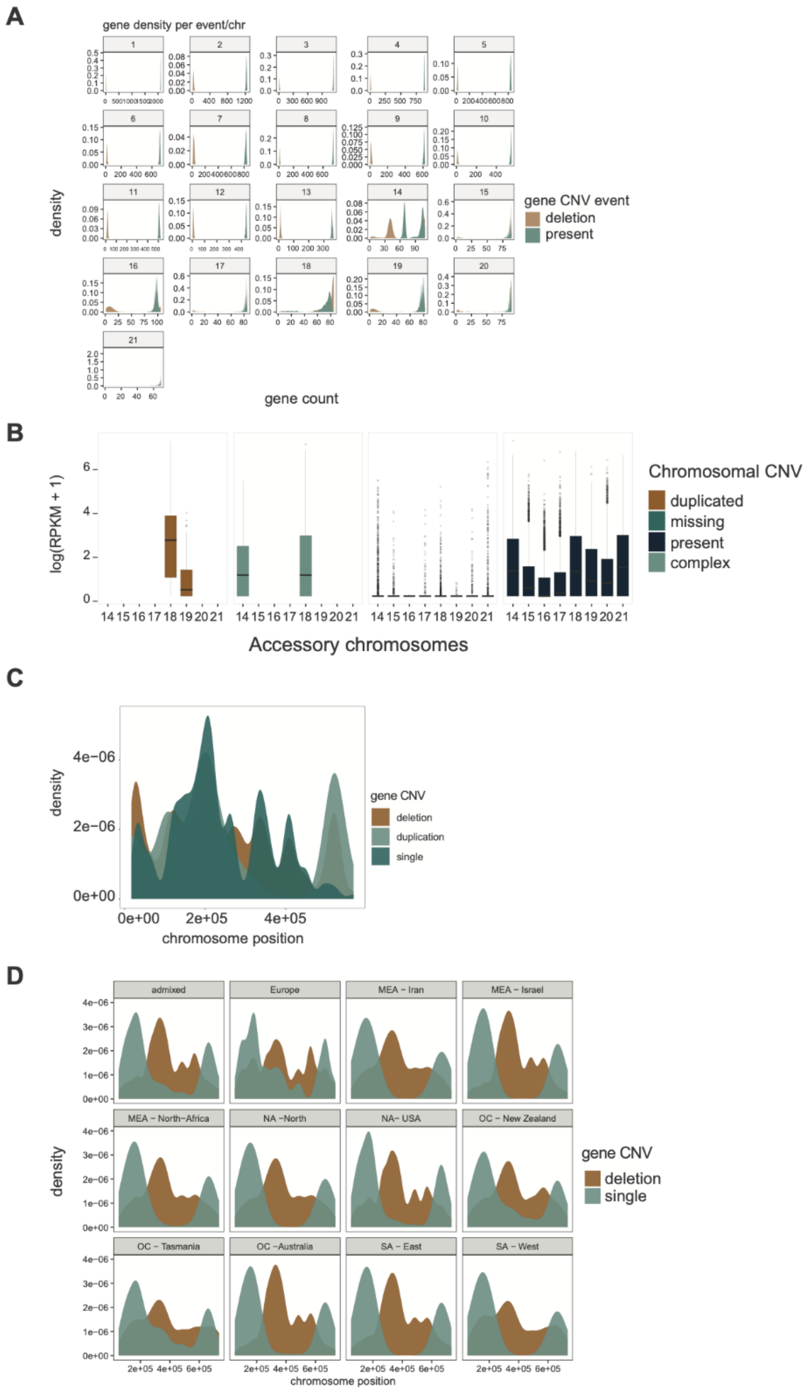


**Supplementary Figure 2.** A) Gene presence/absence density per chromosome in the global genome panel. B) Transcriptional activity assessment under *in vitro* conditions on a subset of the global panel (n=74 strains) Abraham et al. (2023). C) Gene CNV event densities across the chromosome 18 arm. D) Chromosome 14 insertion frequencies across populations.

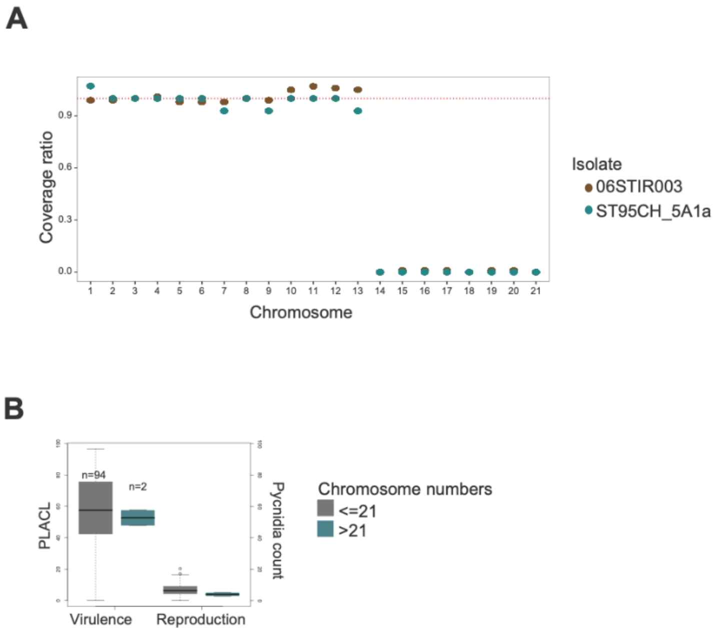


**Supplementary Figure 3.** A) Genome coverage ratio per chromosome of strains carrying solely core chromosomes. B) Virulence (Percent Leaf Area Covered by Lesions, PLACL) and reproduction (pycnidia count) on the wheat host of strains carrying variable total chromosome numbers (Singh et al. 2021). PLACL stands for percentage of leaf area covered by lesions.

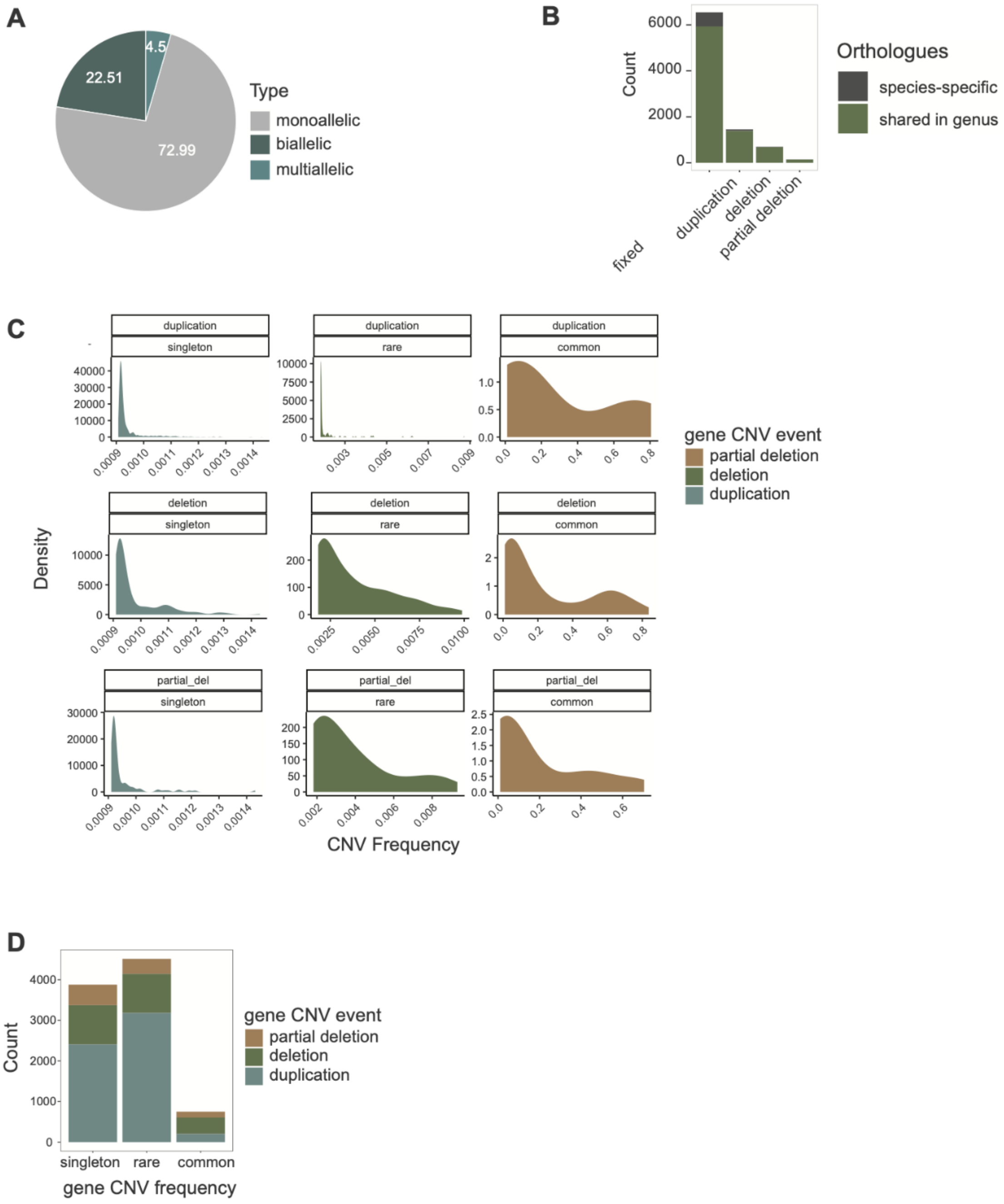


**Supplementary Figure 4.** A) Veen diagram showing the overall percentage of alternative alleles (*i.e.* deletion or duplication) in the global genome panel. Multiallelic refers to genes showing multiple types of events. B) Distribution of orthologs shared between sister species and species-specific genes (*i.e.* unique to *Z. tritici*). C) Density plot showing CNV event distribution of each CNV type. D) Gene CNV events for each CNV frequency type in the unfiltered CNV call dataset.

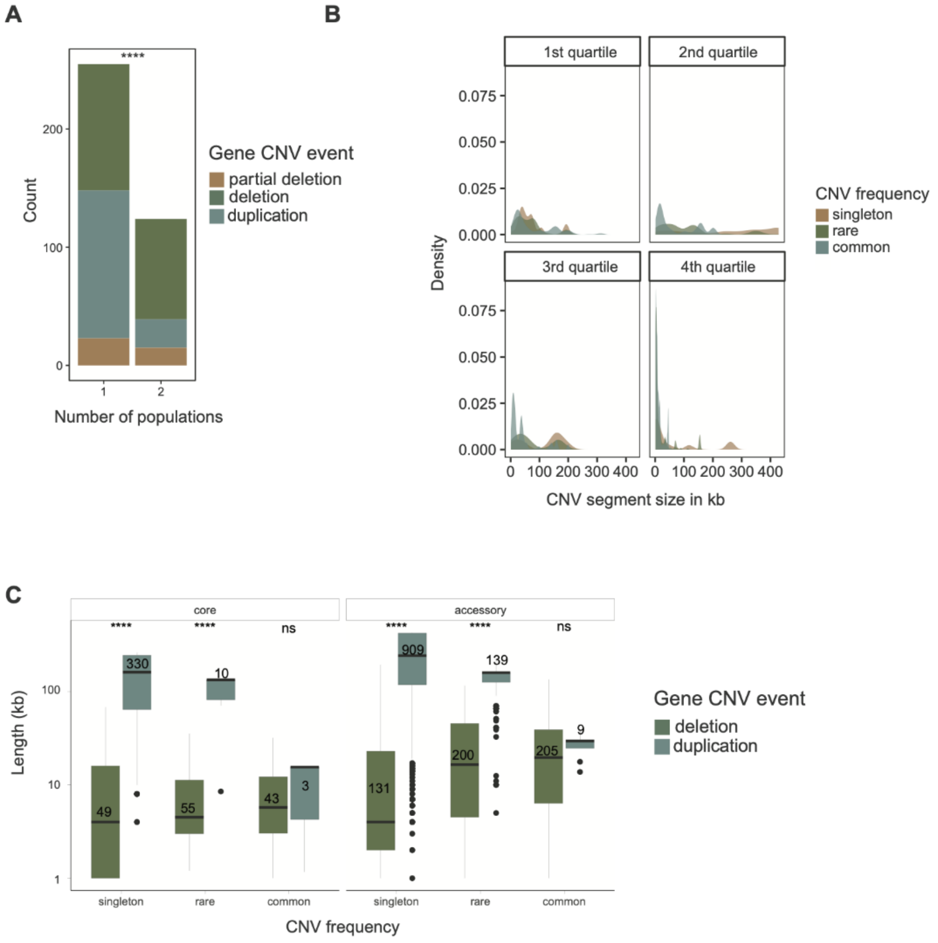


**Supplementary Figure 5.** A) Odds ratio of shared rare frequency gene deletions and duplication in single a population versus population pairs (**** $p$-value < 0.0001). B) CNV segment size variation in the global collection shown separately for each quartile of the CNV segment quality score (QA). C) Overall CNV segment size variation across CNV frequency and CNV event categories in core and accessory chromosomes.

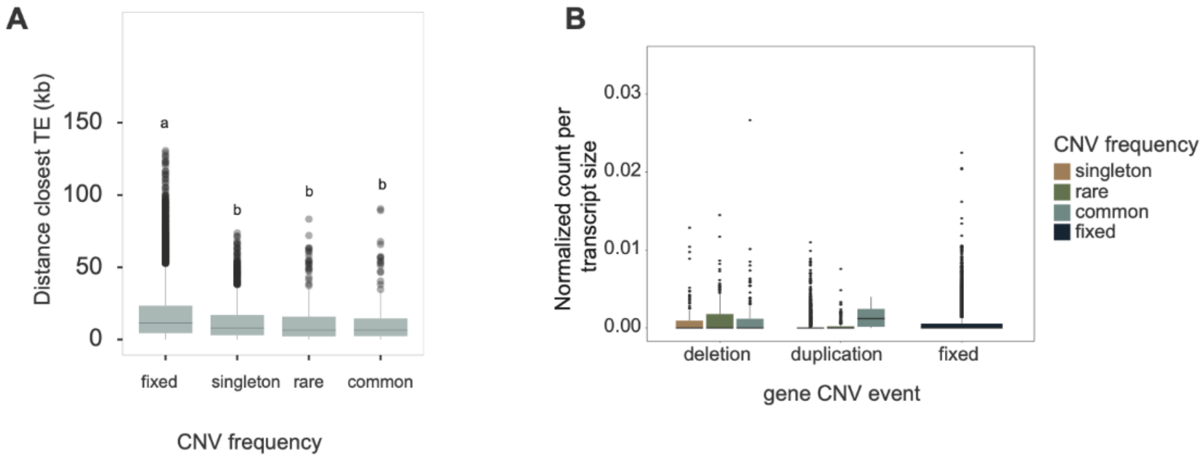


**Supplementary Figure 6.** A) Distance to the closest TE for each gene CNV frequency category. Letters indicate significant differences ($p < 0.05$). B) Predicted protein high-impact SNV variants across gene CNV event categories. Values were normalized by transcript length.

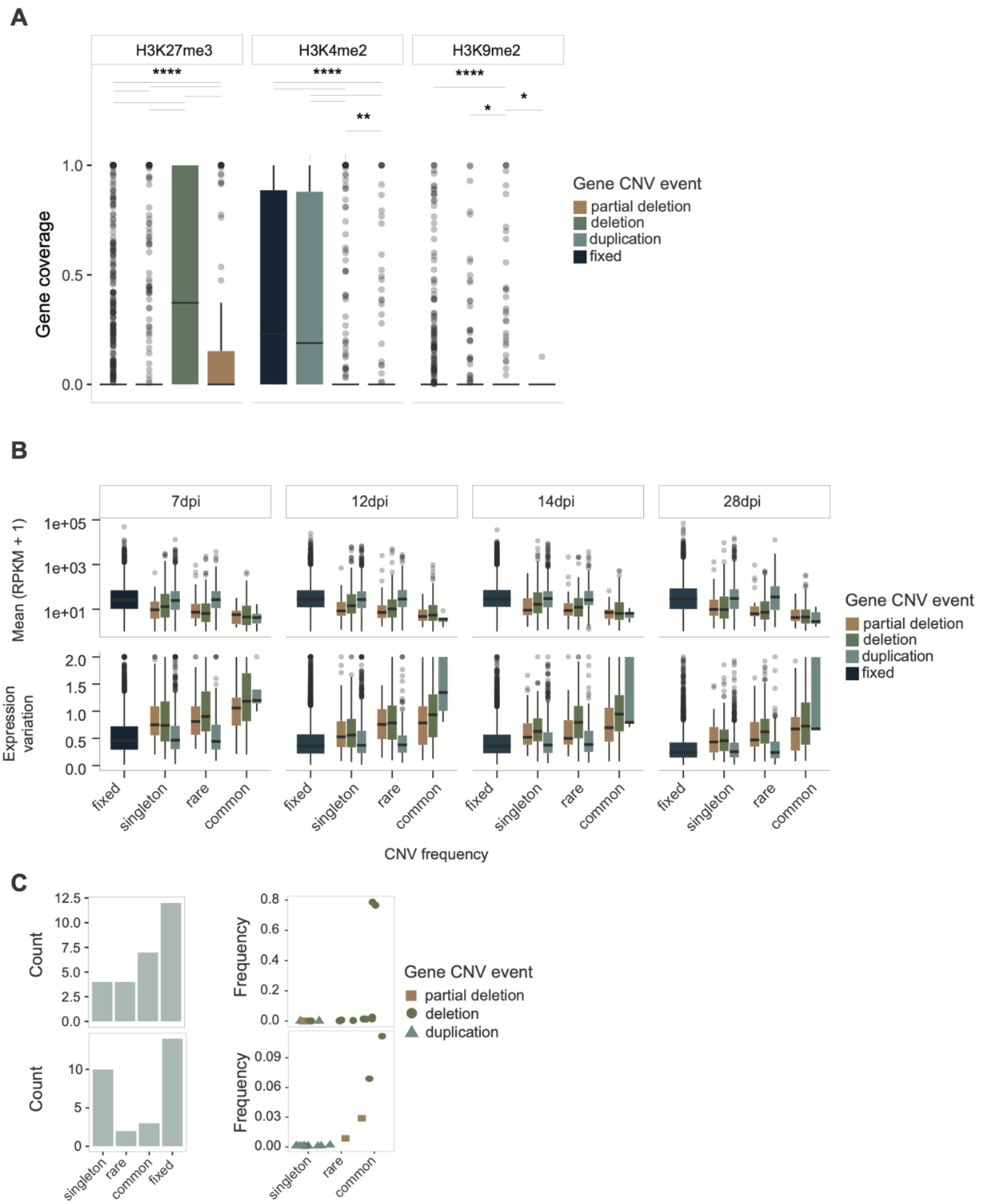


**Supplementary Figure 7**. A) Gene coverage distribution of histone H3K27me3, H3K4me2 and H3K9me2 methylation marks for gene CNV events. B) Gene expression analysis and expression variation during a host infection cycle (7,12,14 and 28 days after infection) across gene CNV events and frequency categories. C) Gene CNV event profile of the most significantly enriched gene ontology (GO) term for biological process (plot above, secondary metabolic process) and molecular function (plot below, serine endopeptidase activity).

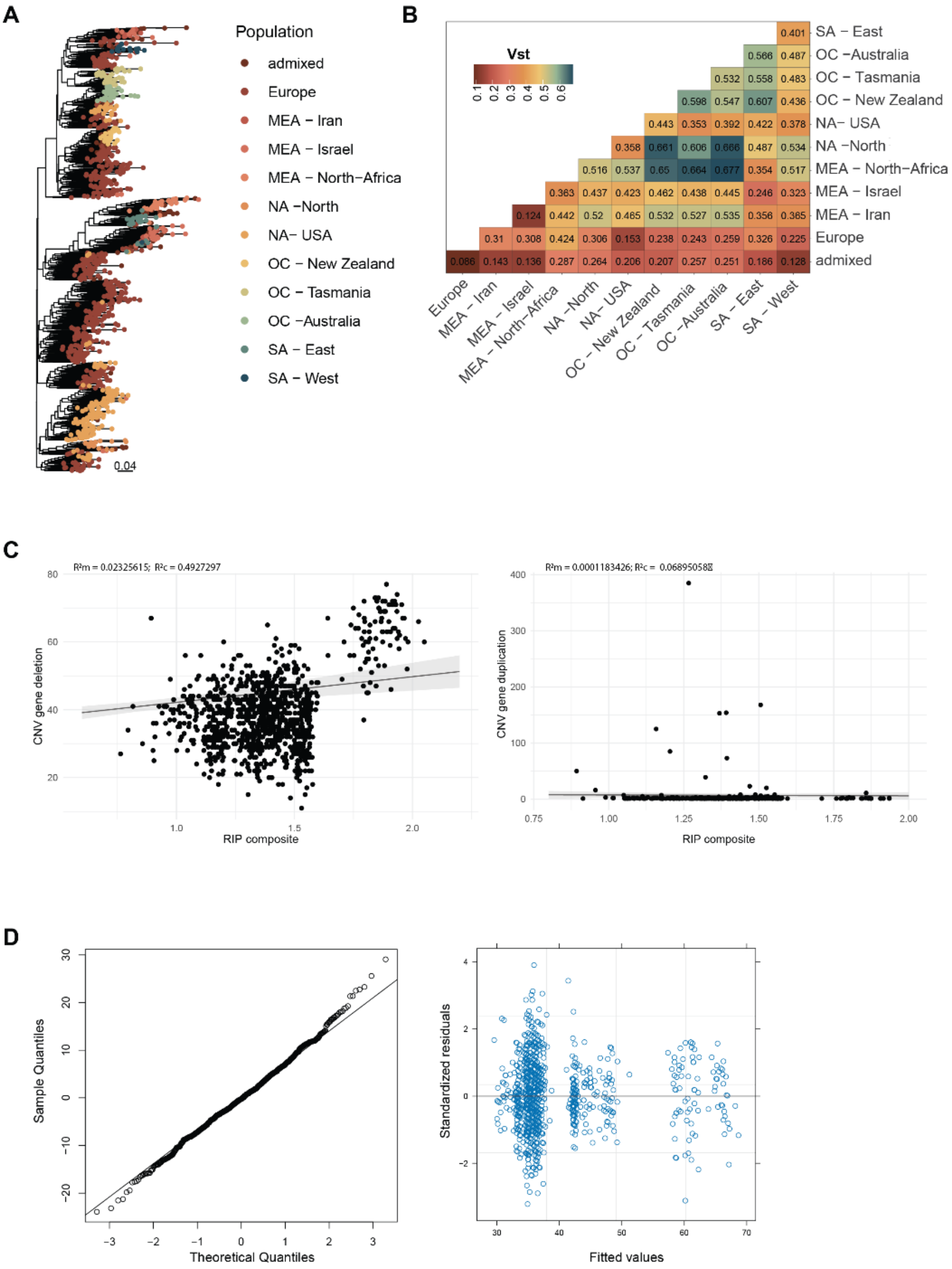


**Supplementary Figure 8.** A) Neighbor-joining tree based on 136 CNV genes filtered for minor allele frequency >0.05 and based on core chromosome genes. The color scheme identifies genetic clusters inferred by SNV analyses. B) CNV-based population differentiation fixation index $V_{ST}$. C) Scatter plot of RIP composite index and gene CNV events (left panel: duplications, right panel: deletions). Fitted line of the fixed effect of the predictor variable (RIP composite index) on the response variable (gene CNVs) accounting for the random effect variables (RIP composite | Population) based on a linear mixed model. $R^2c$ and $R^2m$ refer to the conditional and marginal coefficient of determination for the generalized mixed-effect models, respectively. (D) Quantile-Quantile (QQ) plot of residuals and residual plot of the linear mixed model, indicating the distribution of residuals (vertical axis) against the fitted values (horizontal axis) for the gene deletion model.

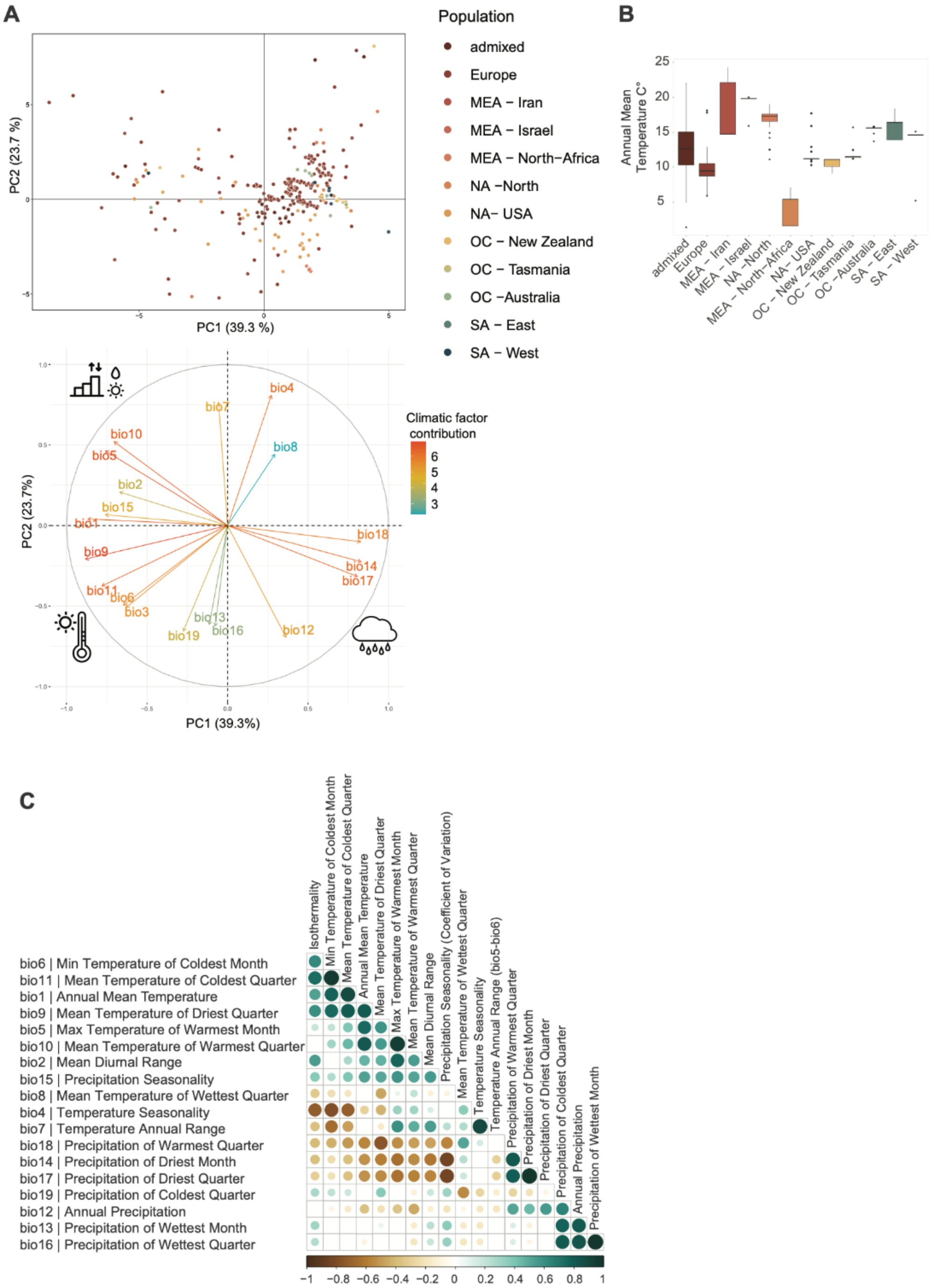


**Supplementary Figure 9.** A) Plot of the first and second principal component based on 19 climatic factors assessed for the geographic location of each sampling location. The color scheme identifies genetic clusters inferred by SNV analyses. The plot below refers to climatic variable contributions to the first and second principal component. Lower-right, lower-left and upper-left areas of the plot represent overall precipitation, temperature and climate range variation variables, respectively. B) Annual mean temperature (bio1)

variation between genetic populations. C) Correlation plot between the 19 climatic factors used for genotype-environment association (*p*-value <0.0001).

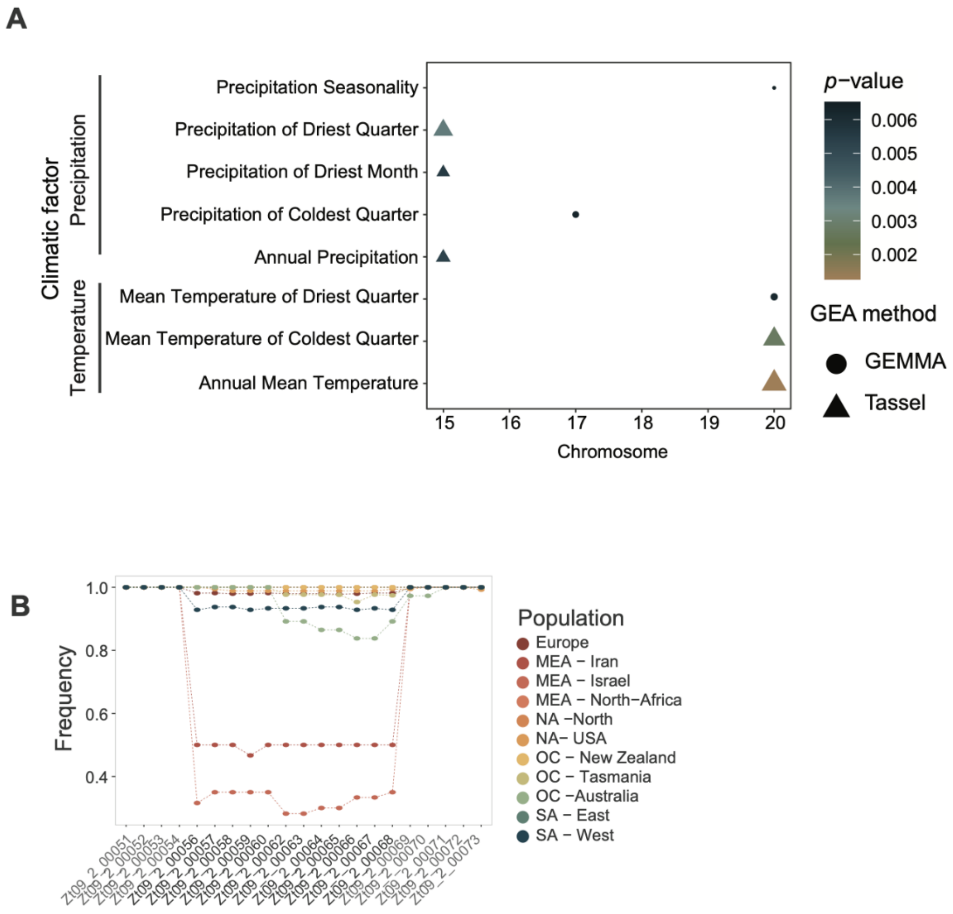


**Supplementary Figure 10.** A) Significant associations from genome-environment association (GEA) analyses (Bonferroni alpha = 0.05) based on chromosome presence/absence variation for 19 climatic factors assessed for the geographic location of each sampling location using two mixed model methods (implemented in Tassel and GEMMA, respectively). B) BGC19 frequency across the genetic clusters. Genes in grey refer to the flanking regions.

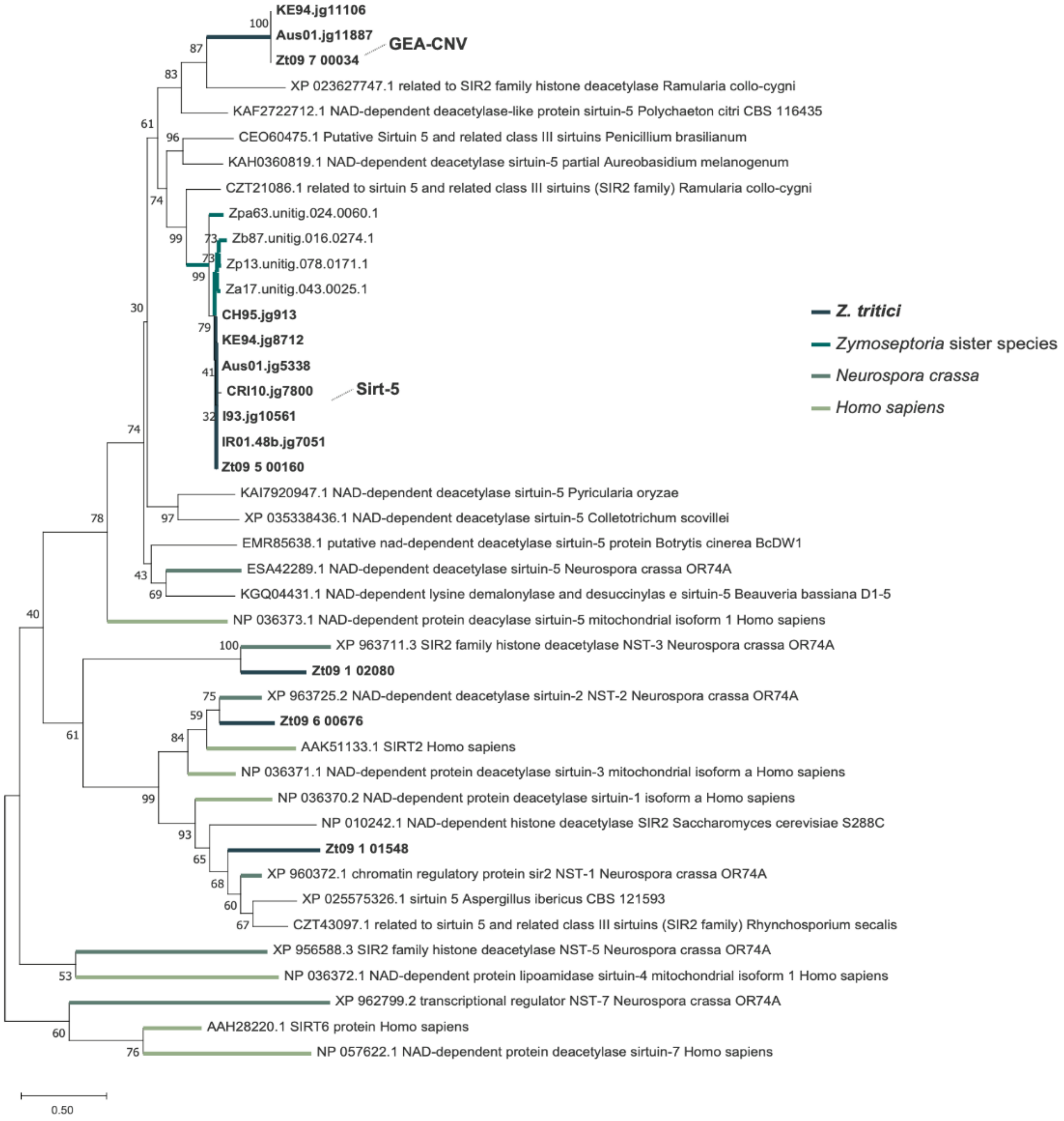


**Supplementary Figure 11.** Evolutionary history of the *Sirtuin* gene family. Unrooted phylogenetic tree for *Sirtuin* orthologs based on maximum likelihood with 1000 bootstrap replicates. The colored line refers to the species.

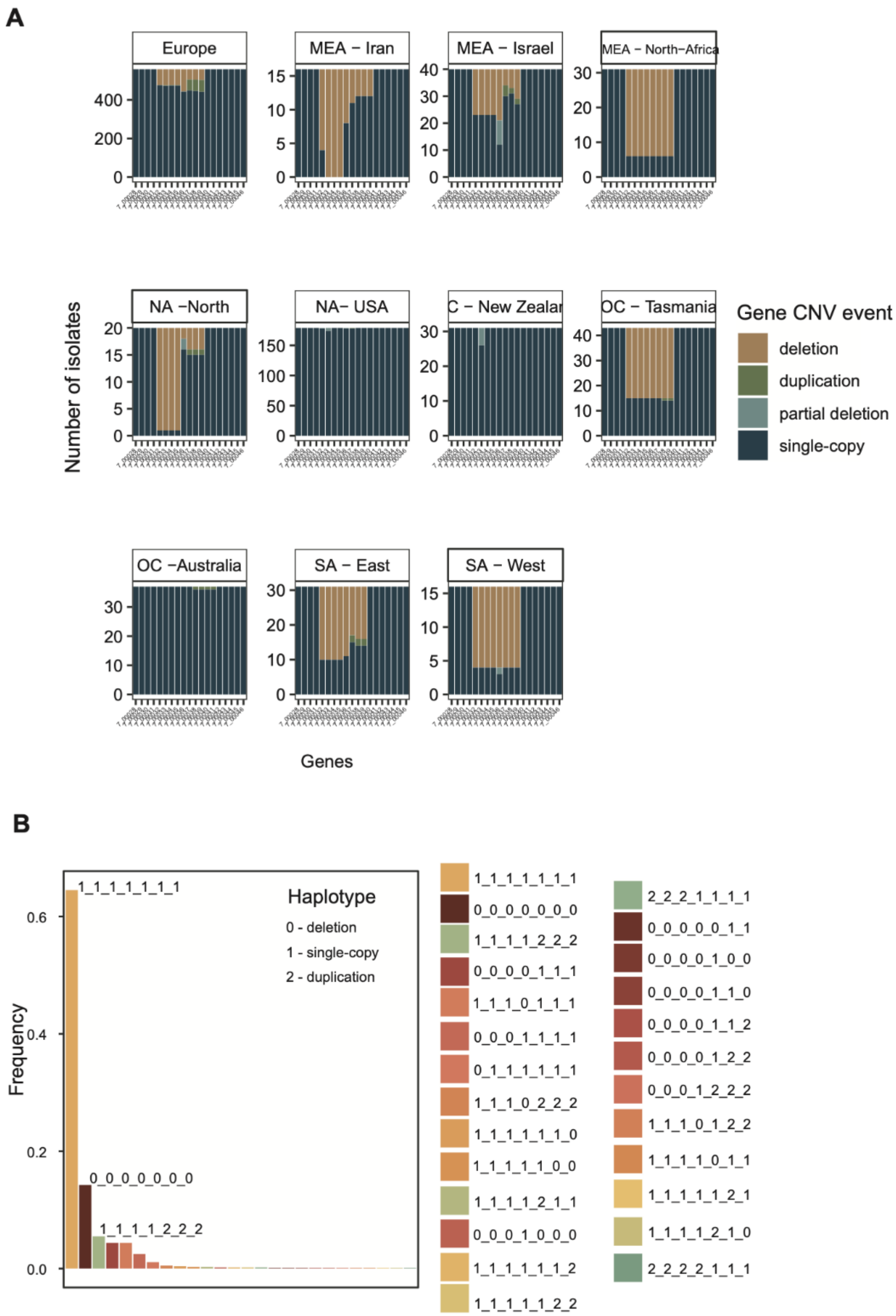


**Supplementary Figure 12.** A) Gene frequency of the *Starship* region across isolates from different genetic clusters. B) *Starship* haplotype frequency based on the unfiltered gene CNV dataset. Haplotypes were ordered by decreasing frequency in the global genome panel.

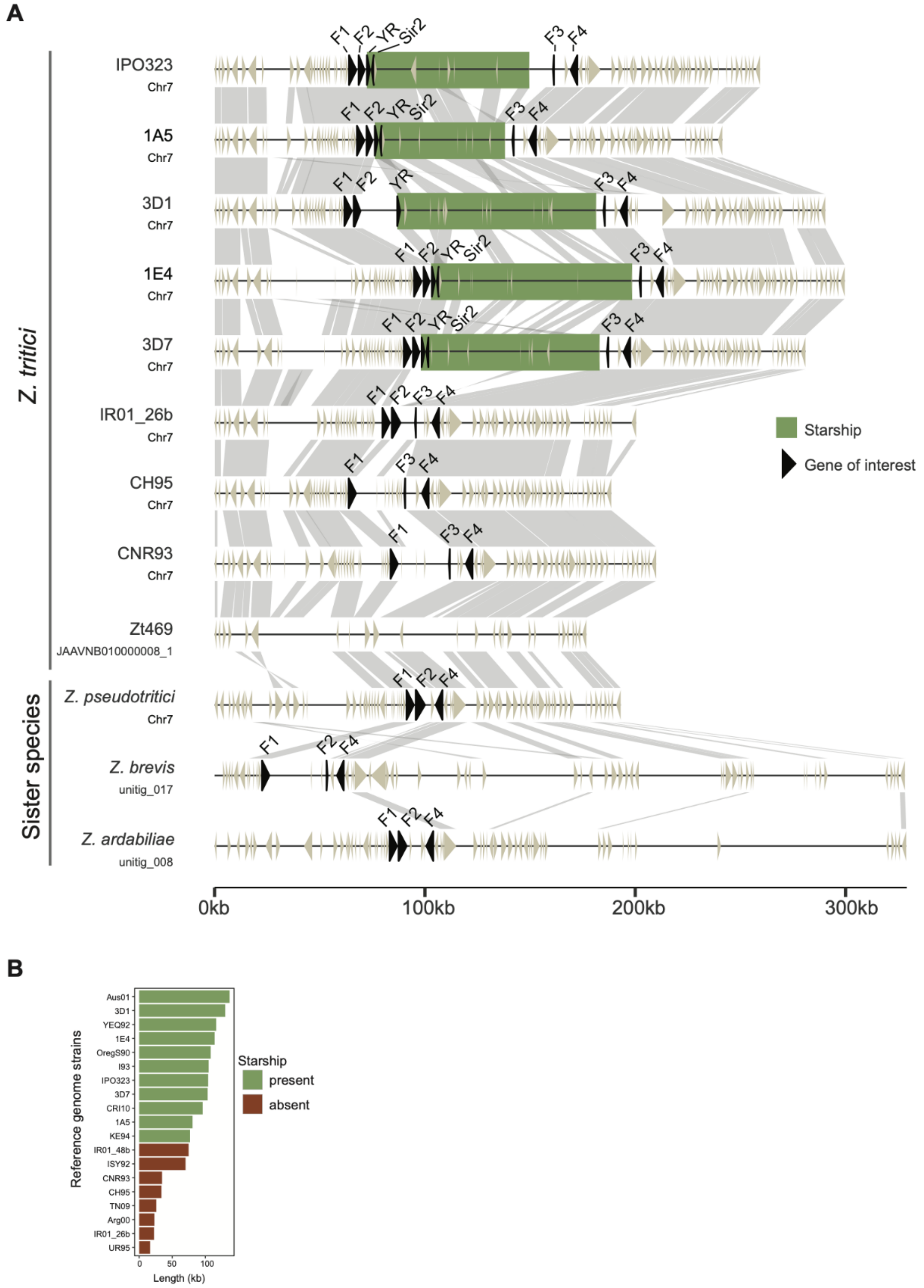


**Supplementary Figure 13.** A) Genome level synteny plot of the *Starship* region between chromosome-level assemblies of *Z. tritici* and sister species strains. Only alignments ≥1000bp and 90% identity are shown. Predicted genes are displayed as arrows and genes of interest are filled in black, including four genes belonging to ortholog groups that have conserved positions flanking the element. YR: tyrosine recombinase (Zt_7_00033); F1: Zt09_7_00031; F2: Zt09_7_00032; F3: Zt09_7_00040; F4: Zt09_7_00042. The green bar identifies the *Starship*. B) Size between flanking region (Zt09_7_00031 up to Zt09_7_00041) of the *Starship* region. Colors refers to presence or absence of the mobile element.

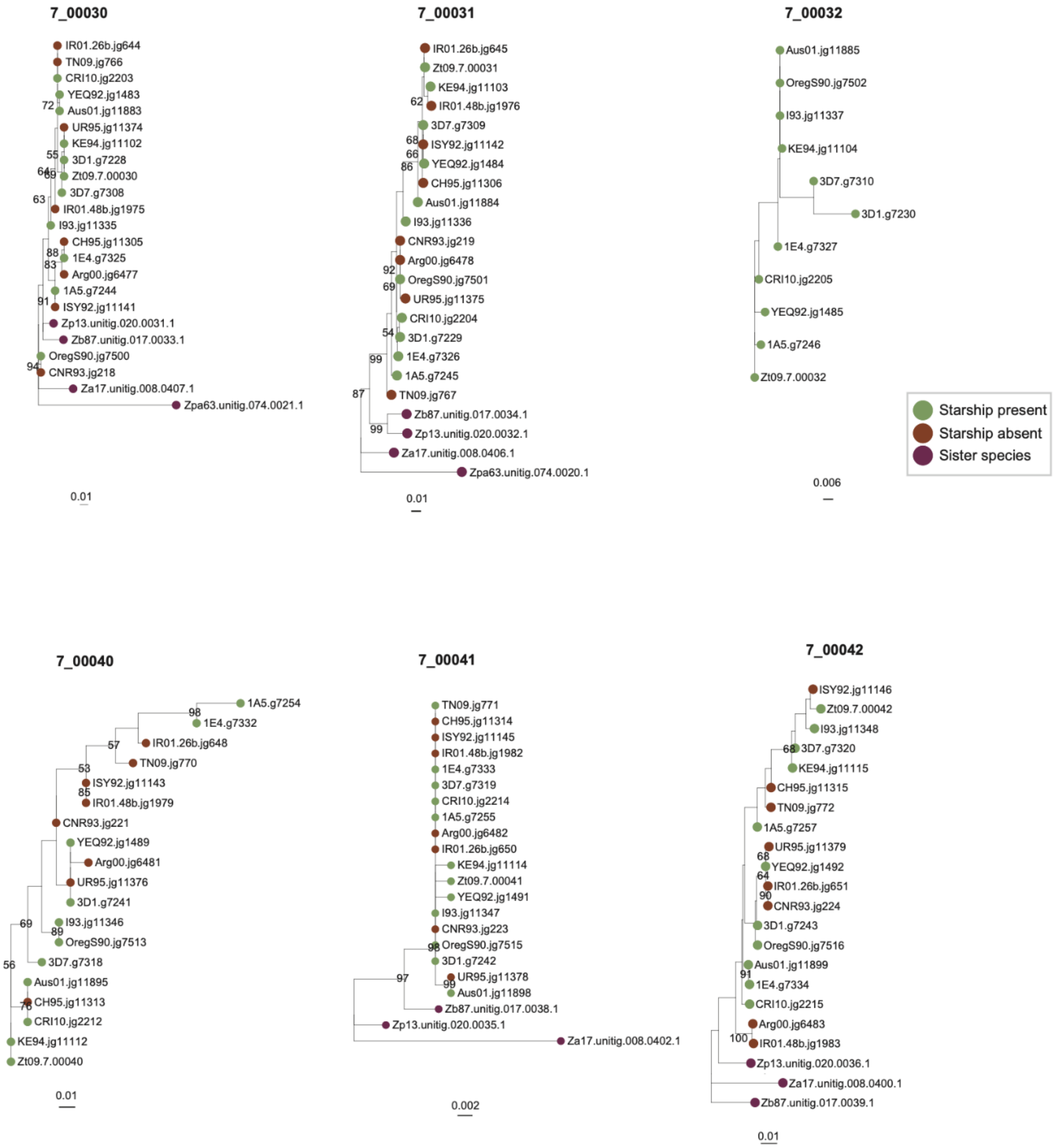


**Supplementary Figure 14.** Phylogenetic trees of the chromosome-level assemblies and sister species protein sequence orthologues flanking the *Starship* region. Trees were constructed using maximum likelihood with a 1000 bootstrap replicates. Color circles identify the presence or absence of the *Starship* across genomes.

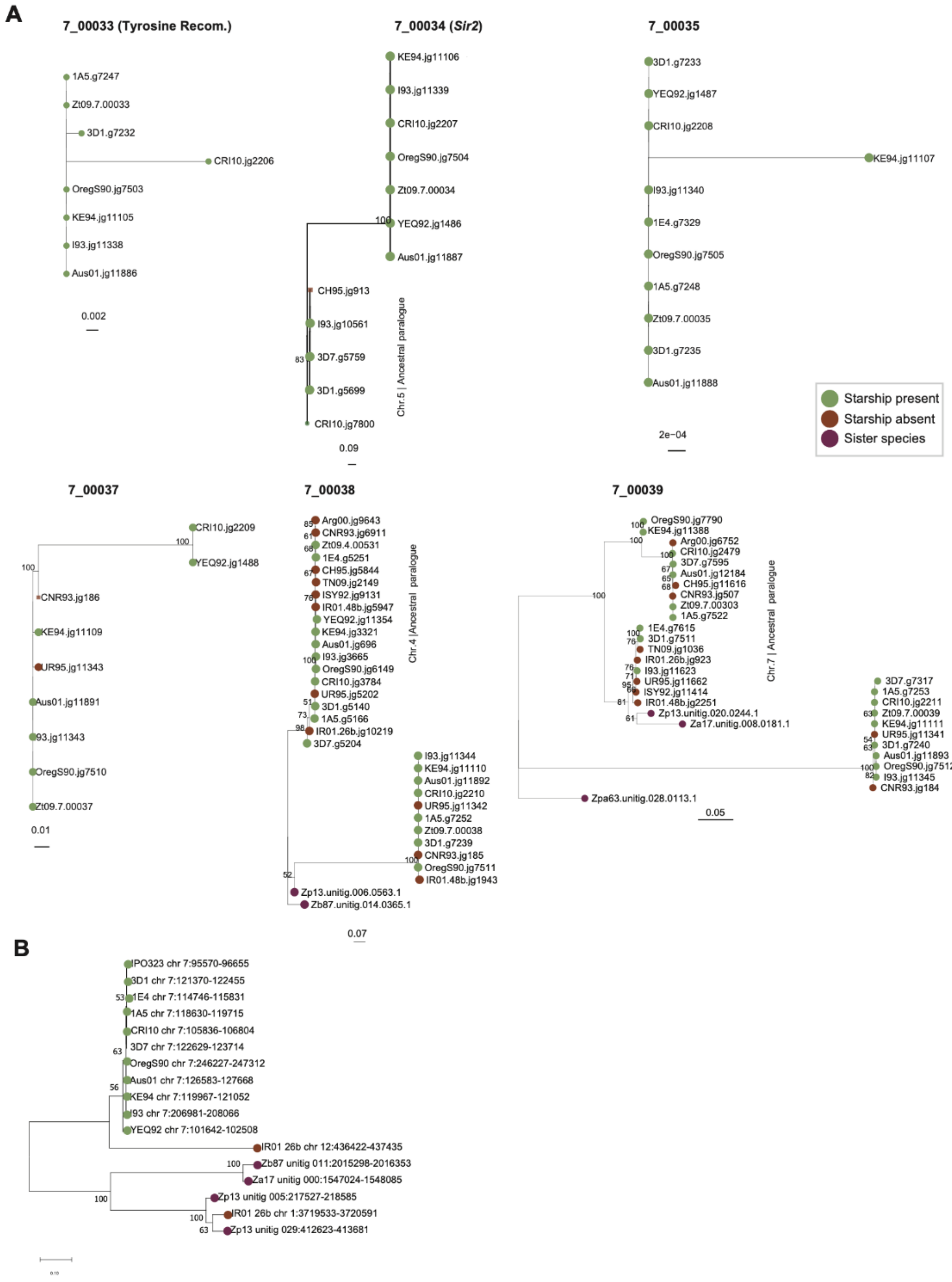


**Supplementary Figure 15.** A) Phylogenetic trees of the chromosome-level assemblies of *Z. tritici* and sister species protein sequence orthologues within the *Starship* region. Trees were constructed using maximum likelihood with 1000 bootstrap replicates. Color circles refer to presence or absence of the *Starship* across genomes. Paralogues in the tree are highlighted. Names identify strains carrying an orthologue. B) Unrooted phylogenetic tree of the tyrosine recombinase gene based on tblastn analysis and built with maximum likelihood and 1000 bootstrap replicates. Names identify chromosome-level assembly strains and the loci coordinates.

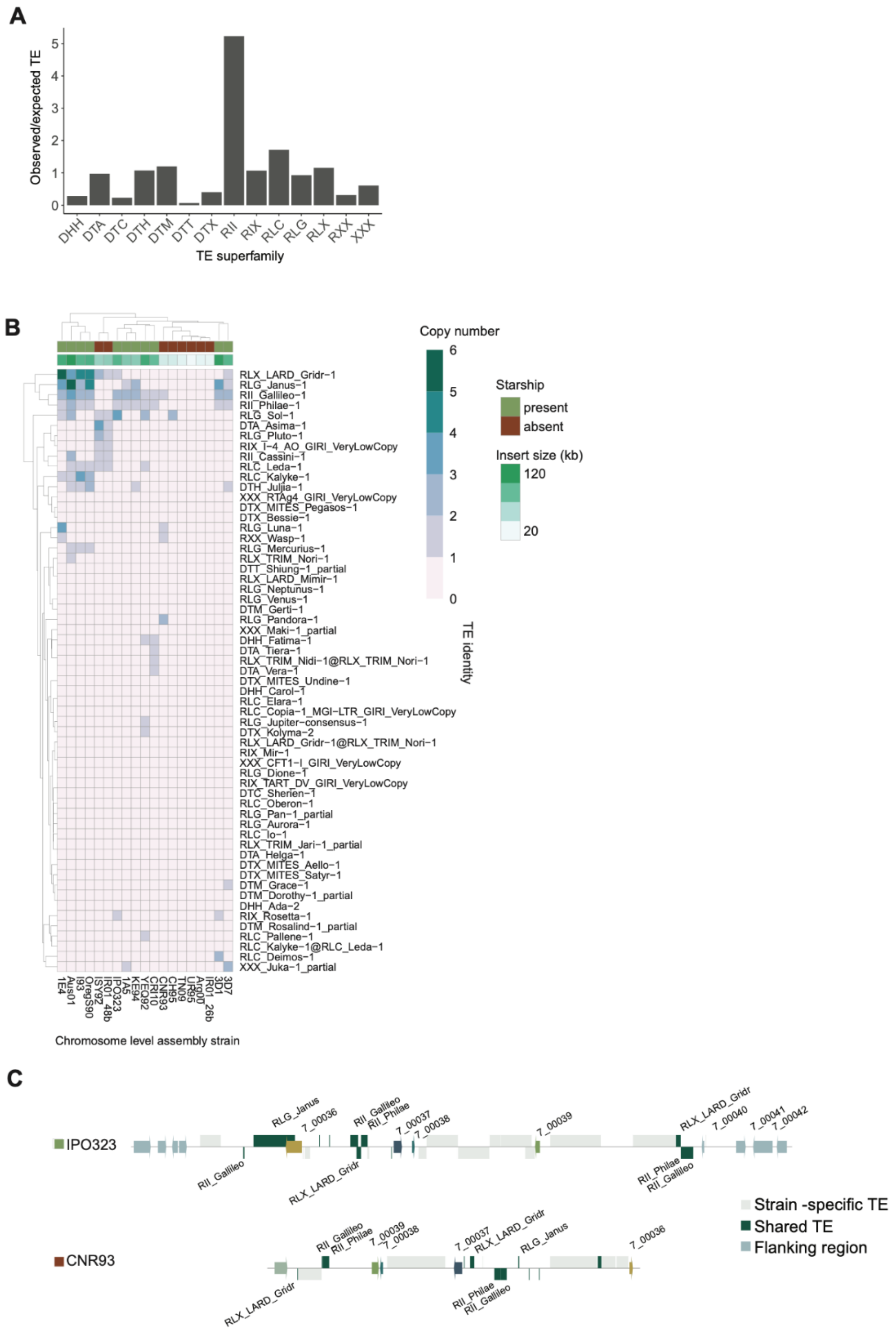


**Supplementary Figure 16.** A) Barplot showing the overall ratio between the *Starship* region compared to genome-wide transposable element content (core chromosomes) in fully assembled genomes. Heatmap of transposable elements within the *Starship* of each chromosome-level assembly. Transposable element annotation was retrieved from Badet 2020 et al. C) Shared TEs between the *Starship* in the IPO323 reference genome and the genomic context in the CNR93 chromosome-level assembly. Arrows represent genes. Blocks represent TEs. @ refers to nested TEs.

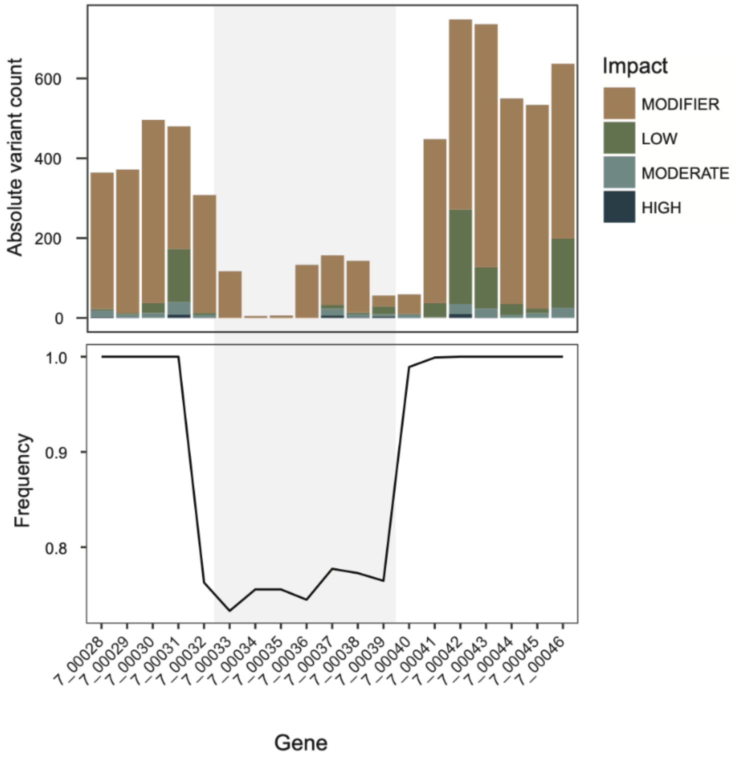


**Supplementary Figure 17.** Absolute variant impact count and loci frequency within the *Starship* region assessed for the global genome panel (n=1104). Grey shading identifies *Starship* cargo genes.
